# Cardiac MAO-A inhibition protects against catecholamine-induced ventricular arrhythmias via enhanced diastolic calcium control

**DOI:** 10.1101/2022.09.29.510139

**Authors:** Qian Shi, Hamza Malik, Jennifer Streeter, Jinxi Wang, Ran Huo, Rachel M. Crawford, Jean C. Shih, Biyi Chen, Duane Hall, E. Dale Abel, Long-Sheng Song, Ethan J. Anderson

## Abstract

**Background:** People with clinical depression exhibit increased risk for cardiac arrhythmias that could be related to differences in catecholamine metabolism. Emerging studies have implicated a pathophysiologic role for monoamine oxidase (MAO-A), which catalyzes catecholamine metabolism in the heart. MAO-A is the pharmacological target of some classes of anti-depressants. Here, we investigated the relationship between MAO-A activity and arrhythmogenesis.

**Methods & Results:** TriNetX database analysis of adult patients with depression (n=11,533) revealed that MAO inhibitor (MAOI) treatment is associated with significantly lower risk of arrhythmias compared with selective serotonin reuptake inhibitor (SSRI) treatment (16.7% vs 18.6%, p=0.0002). To determine a mechanistic link between MAO activity and arrhythmia, we utilized a genetically modified mouse model with cardiomyocyte-specific MAO-A deficiency (cMAO-A^def^). Compared with wild-type (WT) mice, cMAO-A^def^ mice had a significant reduction in the incidence (38.9% vs. 77.8%, p=0.0409) and duration (55.33 ± 26.21s vs.163.1 ± 56.38s, p=0.0360) of catecholamine stress-induced ventricular tachyarrhythmias (VT). Reduced VT risk and duration were associated with altered cardiomyocyte Ca^2+^ handling in the cMAO-A^def^ hearts, including a marked increase in Ca^2+^ reuptake rate, decreased diastolic Ca^2+^ levels, decreased SR Ca^2+^ load and reduced Ca^2+^ spark activity following catecholamine stimulation relative to WT. Further analysis of molecular mechanisms revealed that altered Ca^2+^ handling in the cMAO-A^def^ hearts was related to decreased catecholamine-induced phosphorylation of Ca^2+^/calmodulin-dependent kinase II (CaMKII) and ryanodine receptor 2 (RyR2), and increased phosphorylation of phospholamban (PLB).

**Conclusions:** These findings suggest that MAO-A inhibition in cardiomyocytes mitigates arrhythmogenesis via enhanced Ca^2+^ reuptake that lowers diastolic Ca^2+^ levels thereby diminishing arrhythmic triggers following catecholamine stimulation. Thus, cardiac MAO-A represents a potential target for antiarrhythmic therapy.

---

Depression is a global health crisis, affecting more than 280 million people, or 5% of the global population^1^. Epidemiological studies have found that depression is associated with a substantially increased risk of heart rhythm disorders including atrial fibrillation and ventricular arrhythmia^2–4^. Despite enormous investments in developing new antidepressant drugs, agents that enhance neurotransmission of serotoninergic and catecholaminergic systems have remained the two major pharmacodynamic principles of current drug treatments for depression. Considering heightened catecholaminergic stress is a major pathogenic factor underlying arrhythmogenesis, there is a great need for evaluating the association and mechanisms of antidepressants and the risk of a potentially dangerous heart rhythm disturbances.

Monoamine oxidases (MAOs), the major enzymes responsible for catecholamine metabolism, have been commonly used pharmacological targets since the 1950’s and are still used in treating some patients with depression today^5^. MAO has two isoforms, MAO-A and MAO-B, which are mitochondrial located flavoenzymes that catalyze the oxidative deamination of catecholamines and biogenic amines, producing aldehyde and hydrogen peroxide^6^. Of interest, MAO-A is present in the myocardium of several species including human and rodents. MAO-A principally (but not exclusively) catabolizes serotonin, norepinephrine, and epinephrine, all of which have major pathophysiological implications in the heart including modulation of cardiac inotropy, development of heart failure, arrhythmogenesis, and others. Experimental evidence has revealed that MAO-A is induced and becomes an important source of ROS that contributes to the pathogenesis of heart failure, myocardial ischemia and reperfusion injury, and diabetic cardiomyopathy^7–12^. Recently, MAO-A was reported to regulate inotropic responses in cardiomyocytes by limiting catecholamine accessibility to activate an intracellular pool of β-adrenergic receptors^13,14^.

Despite these prior reports, no cause-effect relationship between cardiac MAO-A and arrhythmogenesis has ever been established. To address this question, we sought to determine whether MAO-A is a viable target for arrhythmogenesis. We provide clinical evidence that patients with clinical depression have a lower incidence of arrhythmic events when treated with MAO inhibitors relative to those treated with selective serotonin reuptake inhibitors. Using a mouse model of cardiomyocyte specific MAO-A inhibition (cMAO-A^def^), we demonstrate that mice deficient in cardiac MAO-A have reduced arrhythmic incidence and duration in response to *in vivo* catecholamine stress, that is associated with faster Ca^2+^ reuptake, lower diastolic Ca^2+^ levels and reduced Ca^2+^ spark activity. Biochemical assays point to altered phosphorylation of important Ca^2+^ regulatory proteins as the molecular basis of improved Ca^2+^ handling and reduced arrhythmogenesis in catecholamine stimulated cMAO-A^def^ hearts. Together, our findings suggest a translational potential of cardiac specific inhibition of MAO-A for the prevention and treatment of cardiac tachyarrhythmias.

## METHODS

### TRINETX STUDY DESIGN AND DATA ANALYSIS

The data used in this study was collected on May 10^th^, 2022, from the TriNetX research network in the United States (Cambridge, MA), which provided access to comprehensive electronic medical records from approximately 105 million patients from 66 healthcare organizations (HCOs). TriNetX is compliant with all data privacy regulations applicable to the contributing HCOs including the Health Insurance Portability and Accountability Act (HIPAA). Any patient level data provided in a data set generated by the TriNetX platform only contains de-identified data as defined in Section §164.514(a) of the HIPAA Privacy Rule. Because this study used only de-identified patient records and did not involve the collection, use, or transmittal of individually identifiable data, this study was exempted from Institutional Review Board approval. <u>Study population:</u> the study population consisted of adults (age 18 to 80) with either depressive episode (ICD10CM:F32) or major depressive disorder, recurrent (ICD10CM:F33) as defined by the International Classification of Diseases (ICD), 10th Revision. Individuals within this study population prescribed with monoamine oxidase inhibitors or selective serotonin reuptake inhibitors were assigned to MAOI and SSRI groups, respectively. Individuals with a diagnosis of attention-deficit hyperactivity disorders (ADHD) (ICD10CM:F90) were excluded to reduce confounding risks. <u>Assessed outcomes:</u> the outcome of interest in this study was the risk of adverse arrhythmic events, including atrial fibrillation and flutter (ICD10CM:I48) and other cardiac arrhythmias (ICD10CM:I49).

### ANIMALS

MAO-A flox/flox (MAO-A ^f/f^) that were used for this study have been previously described^15^. αMyHC-Cre mice expressing Cre recombinase in cardiomyocytes were purchased from Jackson Laboratory (Strain #:011038). MAO-A ^f/f^ mice were bred with αMyHC-Cre expressing mice to generate experimental cohorts including homozygous cardiac-specific MAO-A deficient mice (cMAO-A^def^, MAO-A^f/f^/CRE^+^) and wild-type controls (WT, MAO-A^f/f^/CRE^-^). Both male and female mice at 2-4 months old were randomized and used in this study.

### MAO-A ACTIVITY ASSAY

Cardiomyocyte MAO activity was assessed in 20 μg cell lysate using Amplex™ Red Monoamine Oxidase Assay Kit (ThermoFisher, A12214). p-tyramine is used as the substrate. MAO-A activity is identified by applying specific MAO-A inhibitor clorgyline according to the manufacturer’s directions.

### ELECTROCARDIOGRAPHIC RECORDING AND INDUCTION OF VENTRICULAR ARRHYTHMIAS IN ANAESTHETIZED MICE

Mice were placed on a heating pad (37°C)) and lightly anaesthetized with isoflurane (1-1.5%) in 100% O_2_. Surface electrocardiographs (ECGs) were recorded by PowerLab 8/35 (AD Instruments, Sydney, Australia). Baseline ECGs were recorded for 5 min when stable heart rates were reached, followed by an additional 30 min recording following intraperitoneal (i.p.) administration of epinephrine (adrenaline) (2 mg/kg) and caffeine (120 mg/kg)^16,17^. Baseline parameters including heart rate, P-wave duration, PR-interval, QRS-interval, and QT-interval were obtained via the ECG analysis module of LabChart 8 (AD Instruments, Sydney, Australia). Analysis of cardiac arrhythmias was performed manually. Definitions of ventricular arrhythmias were based on the Lambeth Conventions II^18^. Both non-sustained ventricular tachycardia (VT; run of 4-10 consecutive single ventricular ectopic beats) and sustained VT (run of 10 or more consecutive single ventricular ectopic beats) were included in the calculation of VT incidence and duration.

### CONFOCAL CA^2+^ IMAGING IN INTACT MOUSE HEARTS

Excised hearts were loaded with Rhod-2 AM (rhodamine 2 acetoxymethyl ester; 5μM, AAT Bioquest) in Krebs–Henseleit (KH) solution (120 mM NaCl, 24 mM NaHCO_3_, 11.1 mM glucose, 5.4 mM KCl, 1 mM MgCl_2_, 0.42 mM NaH_2_PO_4_, 10 mM taurine and 5 mM creatine, oxygenated with 95% O_2_ and 5% CO_2_) at room temperature for 40 min via a retrograde Langendorff perfusion system. Hearts were later transferred to another Langendorff apparatus (37°C) attached to the confocal microscope system after Rhod-2 AM loading was completed. To minimize motion artefacts during Ca^2+^ imaging, blebbistatin (5–10 μM) was added to the perfusion solution. *In situ* confocal line-scan imaging of Ca^2+^ signals arising from epicardial myocytes was performed and acquired at a rate of 3.07 ms/line. Ca^2+^ transients were recorded under electrical pacing at 5 Hz or 50 Hz (by placing a platinum electrode onto the surface of ventricle apex). Analysis of Ca^2+^ imaging data was performed offline using custom-compiled routines in IDL, as previously described^19^.

### ISOLATION OF CARDIOMYOCYTES

Mouse ventricular myocytes were isolated via enzymatic digestion as previously described^20^. Briefly, the hearts were quickly excised and perfused on a Langendorff apparatus at 37 °C with normal Ca^2+^ free Tyrode’s solution (containing the following in mM: NaCl, 137; KCl, 5.4; MgCl_2_, 2.0; NaH_2_PO4, 0.33; D-Glucose, 10.0; and HEPES, 10.0; pH 7.40 at 37 °C). After ∼5 min of perfusion, the perfusate was switched to Tyrode’s solution containing Collagenase (Type 2, 1mg/ml, Worthington) and protease (0.05 mg/ml, Sigma-Aldrich) for the digestion of the connective tissue. After ∼20 min of digestion, single ventricular myocytes were isolated from the dissected and triturated ventricles and stabilized in Tyrode’s solution containing BSA (1%). After gradually reintroducing Ca^2+^, the final Ca^2+^ tolerant cardiomyocytes were resuspended in 1.8 mM Ca^2+^ Tyrode’s solution and maintained at room temperature.

### CONFOCAL CA^2+^ IMAGING IN SINGLE ISOLATED CARDIOMYOCYTES

All Ca^2+^ imaging experiments were performed at 37 °C. After isolation, myocytes were loaded with Rhod 2-AM (5μM, AAT Bioquest) for 25 min at room temperature, followed by additional 25 min of incubation in fresh Tyrode’s solution, to wash out excess dye and allow the complete de-esterification of Rhod 2-AM. Myocytes were then seeded on a laminin coated perfusion chamber, which is mounted on the inverted confocal microscope (ZEISS 510, LSMTech, Pennsylvania, USA) equipped with a 63×, 1.4 NA oil immersion objective. Rhod 2-AM was excited with the 561 nm line of an argon laser and emission was collected by 575 nm longpass filter. To assess Ca^2+^ <u>dynamics in a single cardiomyocyte</u>, myocytes were paced using extracellular platinum electrodes at various frequencies (1, 3, 5Hz). Ca^2+^ transients were recorded accordingly in the line-scan mode along the long axis for 2000 lines at a rate of 1.93ms/line. To examine catecholamine’s effect on myocyte Ca^2+^ handling, same procedures were carried during perfusion with norepinephrine (NE, 1μM). To assess the sarcoplasmic reticulum (SR) Ca^2+^ content, after recording Ca^2+^ transients under 1Hz pacing as described above, electrical stimulation was stopped, and 20 mM caffeine was applied locally to rapidly induce total Ca^2+^ release from the SR. The caffeine induced Ca^2+^ transient amplitude was used as an estimate of SR Ca^2+^ content. The time constants (Tau) of twitch (1Hz pacing) and caffeine-induced Ca^2+^ transient decay were calculated from mono-exponential curve fitting, which reflect the contribution of sarco/endoplasmic reticulum Ca^2+^-ATPase (SERCA) and sodium-calcium exchanger (NCX) to diastolic Ca^2+^ removal, respectively^21,22^. To assess Ca^2+^ <u>spark activity</u>, quiescent cardiomyocytes were recorded in the line-scan mode along the long axis for 1000 lines at a rate of 1.93ms/line. Six consecutive recordings from one quiescent cardiomyocyte were used for Ca^2+^ spark analysis by a custom IDL program as previously described^20^.

### IMMUNOBLOTTING

For protein extraction, frozen heart tissues were collected in Precellys^®^ Tissue Homogenizing Mixed Beads Kit (2.0 mL) and homogenized using a PRECELLYS homogenizer (Bertin) containing ice-cold RIPA lysis buffer (R0278, Sigma-Aldrich), supplemented with protease inhibitor mix (cOmplete, Mini Protease Inhibitor Cocktail, Roche). Solubilized heart homogenates were obtained by centrifugation at 13,200 rpm at 4°C for 30 min. The resulting supernatants were quantified for protein using a Bradford BioRad protein assay. The supernatant was mixed with LDS loading buffer and then resolved in a gradient NuPAGE gel (4%-12%, Invitrogen). NuPAGE gel resolved proteins were transferred to PVDF membranes at 30 V overnight at 4 C. After blocking with TBS containing 0.5 % Tween-20 and 5 % non-fat milk powder, specific proteins were detected with anti-RyR2 (MA3-916, Sigma), anti-Ca_V_1.2 (CACNA1C) (ACC-003, Alomone labs), anti-Na_V_1.5 (SCN5A) (ASC-005, Alomone labs), anti-Phospholemman (PLM, PA5-792881), anti-NCX1 (79350, Cell Signaling), SERCA2a (MA3-919, Thermo Fisher), anti-Calsequestrin (CSQ, PA1-913, Thermo Fisher), anti-RyR2 (pSer2814) (A010-31AP, Badrilla), anti-RyR2 (pSer2030) (A010-32, Badrilla), anti-RyR2 (pSer2808) (A010-30, Badrilla), anti-CaMKII (phospho T286) (ab32678, Abcam), anti-CaMKII (ab181052, Abcam), anti-β2-adrenergic receptor (β2AR) (A-B2AR, Badrilla), anti-β1-adrenergic receptor (β1AR) (ab3442, Abcam), PDE4D (PD4-401AP, FabGennix), GAPDH-HRP (MA5-15738-HRP, Thermo Fisher), anti-Phospholamban (PLB-pSer16) (A010-12, Badrilla), anti-Phospholamban (PLB-pThr17) (A010-13, Badrilla), and anti-Phospholamban (PLB) (A010-14, Badrilla) antibodies. All primary antibodies were revealed with HRP-conjugated goat secondary antibodies using the Bio-Rad ChemiDoc MP and detected using SuperSignal™ West Pico or Femto Chemiluminescence Substrate (Thermo Fisher Scientific). Protein densitometry was analyzed by Image Lab software (6.0).

### STATISTICS

All statistical analyses for the TriNetX study were completed on the TriNetX research platform. Potential confounding factors including patient characteristics (age, sex, race/ethnicity) and comorbidities (diabetes, essential hypertension, hyperlipidemia, opioid use disorder) was considered in this study. Therefore, a 1:1 propensity score technique was used to match cohorts, mitigate risk of bias, and obviate the need for covariate adjustments. The propensity score matching analysis was performed using a multivariable logistic regression model and nearest neighbor algorithms with a tolerance level of 0.01 and difference between propensity scores ≤0.1^23^. Risk difference (RD), risk ratio (RR) and odds ratio (OR) with 95% confidence intervals (CIs) for the arrhythmic outcome was calculated and *p* < 0.05 indicates statistical significance between groups. Data obtained from animal models and related biochemistry studies are presented as mean ± SEM. Statistical differences were determined by GraphPad Prism version 9.0 for Windows (GraphPad Software, Inc). The incidence of VT was compared using Fisher’s exact test. The VT duration was compared with the Mann-Whitney U test as the data were not normally distributed. Normally distributed data were analyzed by parametric tests: one-way analysis of variance (ANOVA) followed by Tukey post hoc for > 2 groups or Student’s t-test for 2 groups. P <0.05 were considered statistically significant.

## RESULTS

### MAOI TREATMENT IS ASSOCIATED WITH LOWER RISK OF ARRHYTHMIC EVENTS IN ADULTS WITH DEPRESSION

Given the known association between depression and risk of arrhythmias, we sought to determine whether arrhythmic events correlated with the class of antidepressant medication used by asking whether MAO-A activity correlates with arrhythmic outcomes. We conducted a retrospective matched cohort comparison of patients with depression, treated either with MAOIs or SSRIs utilizing the TriNetX database and analytics platform. 11,533 individuals were identified who were treated with MAOIs while 2,025,313 individuals were found to have been treated with SSRIs. The baseline characteristics of these two groups varied prior to matching. Specifically, the MAOI group had a higher mean age, percentage of males, Caucasian ethnicity, prevalence of essential hypertension, dyslipidemia, and more likely to be prescribed cardiovascular medications compared with SSRI group. The MAOI group also had decreased prevalence of opioid use-related disorders compared with the SSRI group (Table 1, before propensity score matching). To avoid these confounders, 1:1 propensity score matching was performed, using logistic regression to balance the groups^23^. After matching, both groups were well balanced for all clinical and demographic variables (Table 1, after propensity score matching).

**Table 1.** Baseline Characteristics (Demographics and health) n (%) of Patients with Depression Treated with MAOi or SSRI Before and After Propensity Score Matching^*^

|  | MAOi group<br>(n=11,533) | SSRI group<br>(n=2,025,313) | *p* Value | MAOi group<br>(n=11,533) | SSRI group<br>(n=11,533) | *p* Value |
| --- | --- | --- | --- | --- | --- | --- |
|  | Before propensity score matching |  |  | After propensity score matching |  |  |
| Age, Mean±SD | 63.6±13.5 | 51±17.2 | <0.0001 | 63.6±13.5 | 63.6±13.5 | 0.9511 |
| Sex, *n* (%) |  |  |  |  |  |  |
| Male, | 5,723 (49.627) | 612,085 (30.222) | <0.0001 | 5,723 (49.627) | 5,736 (49.74) | 0.8641 |
| Female | 5,809 (50.373) | 1,412,796 (69.757) | <0.0001 | 5,809 (50.373) | 5,796 (50.26) | 0.8641 |
| Ethnicity, *n* (%) |  |  |  |  |  |  |
| Hispanic or Latino | 478 (4.145) | 167,750 (8.283) | <0.0001 | 478 (4.145) | 491 (4.258) | 0.6696 |
| Race, *n* (%) |  |  |  |  |  |  |
| White | 8,892 (77.107) | 1,410,754 (69.656) | <0.0001 | 8,892 (77.107) | 8,890 (77.09) | 0.9750 |
| Black or African American | 506 (4.388) | 241,335 (11.916) | <0.0001 | 506 (4.388) | 506 (4.388) | 1.0000 |
| Diagnoses, *n* (%) |  |  |  |  |  |  |
| Essential hypertension | 4,132 (35.831) | 625,024 (30.861) | <0.0001 | 4,132 (35.831) | 4,134 (35.848) | 0.9781 |
| Disorders of lipoprotein metabolism | 3,492 (30.281) | 504,799 (24.924) | <0.0001 | 3,492 (30.281) | 3,490 (30.264) | 0.9771 |
| Opioid related disorders | 226 (1.960) | 48,325 (2.386) | 0.0028 | 226 (1.960) | 238 (2.064) | 0.5736 |
| Diabetes mellitus | 1,775 (15.392) | 301,880 (14.905) | 0.1435 | 1,775 (15.392) | 1,742 (15.106) | 0.5455 |
| Alcohol related disorders | 694 (6.018) | 115,019 (5.679) | 0.1169 | 694 (6.018) | 690 (5.983) | 0.9117 |
| Cardiovascular Medications** | 7,886 (68.384) | 1,037,504 (51.227) | <0.0001 | 7,886 (68.384) | 7,877 (68.306) | 0.8986 |

\*, The 1:1 propensity score matching technique was performed using a multivariable logistic regression model to balance the baseline characteristics of the population. \*\*, Cardiovascular medications include beta-blockers, antiarrhythmics, diuretics, lipid lowering agents, antianginals, calcium channel blockers, ACE inhibitors.

As shown in Figure 1A, 1,929 individuals encountered arrhythmic events among the MAOI group (16.726 %, n = 11,533) compared to 2,146 individuals among the SSRI group (18.607%, n = 11,533). The MAOI group also had significantly lower risk (risk difference: −1.882%; 95% CI: −2.866%, −0.897, p=0.0002) of developing arrhythmic events as well as fewer incidences of arrhythmia (mean: 7.721 vs 9.498, p =0.0035, Figure 1B) compared to individuals treated with SSRIs. These data suggest MAOIs may lower the risk of cardiac arrhythmic outcomes in patients with depression relative to those treated with SSRIs.

**Figure 1.**
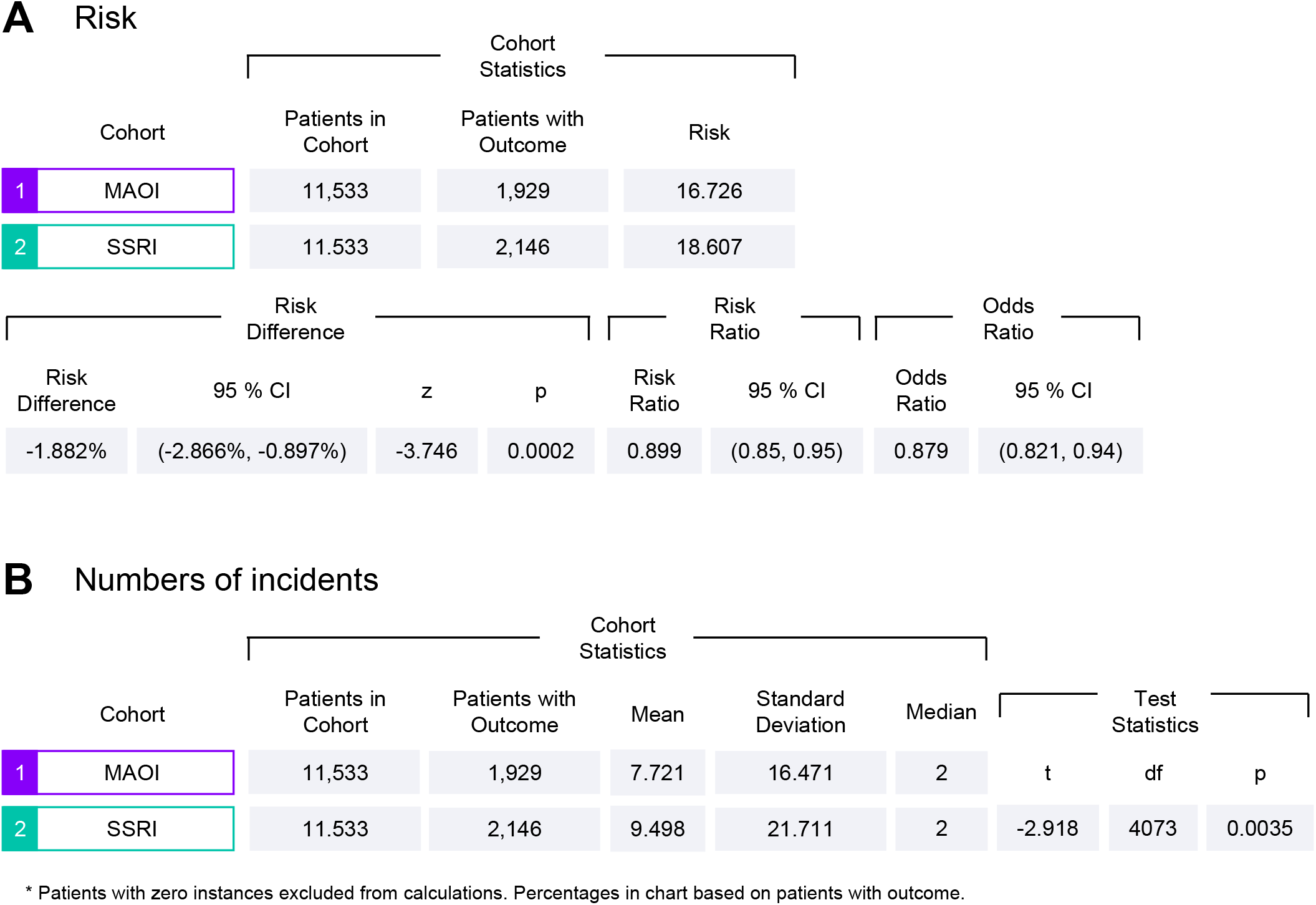
TriNetX analysis of arrhythmia risk among adult patients with depression prescribed either MAO inhibitors (MAOI) or selective serotonin reuptake inhibitors (SSRI)

### MICE WITH CARDIAC MAO-A DEFICIENCY ARE PROTECTED FROM CATECHOLAMINE-INDUCED VENTRICULAR TACHYARRHYTHMIAS

Both MAO-A and MAO-B isoforms are widely expressed in various tissues throughout the body. Although human cardiomyocytes express both, MAO-A appears to be the predominant isoform ^24,25^. To elucidate the link between MAO-A and arrhythmogenesis, we selectively disrupted the *Maoa* gene in cardiomyocytes (cMAO-A^def^) using an αMHC promoter-driven Cre-LoxP system and tested whether cMAO-A^def^ mice would be protected against VT induced by sympathetic system over-drive. Enzyme activity assays confirmed that MAO-A activity was reduced more than 50% in isolated cardiomyocytes (Supplemental Figure 1). Catecholamine stress was applied via a standard protocol through intraperitoneal injection of epinephrine (2 mg/kg)/caffeine (120 mg/kg) to mice under anesthesia^19,26–28^. Incidence and duration of VT were evaluated by ECG recording in both WT and cMAO-A^def^ mice. While both groups have normal sinus rhythm under baseline conditions, catecholamine-inducible VT were significantly more reproducible in WT mice (14 of 18 animals, 78%) compared with cMAO-A^def^ mice (7 of 18 mice, 39%; p=0.0409) (Figure 2A-2C). VT duration was also significantly shorter in cMAO-A^def^ mice compared to WT mice (Figure 2D, 55.33±26.21s vs.163.1±56.38s, p=0.0360). Heart rate and conduction intervals obtained from surface ECG recordings were comparable among WT and cMAO-A^def^ mice both at rest and at the end of 30-minute catecholamine stimulation (Figure 2E and Supplemental Figure 2A-2H). cMAO-A^def^ mice had slightly increased QRS intervals at resting condition (Supplemental Figure 2D) that may be due to modest increases in heart weights (Supplemental Figure 3B).

**Figure 2.**
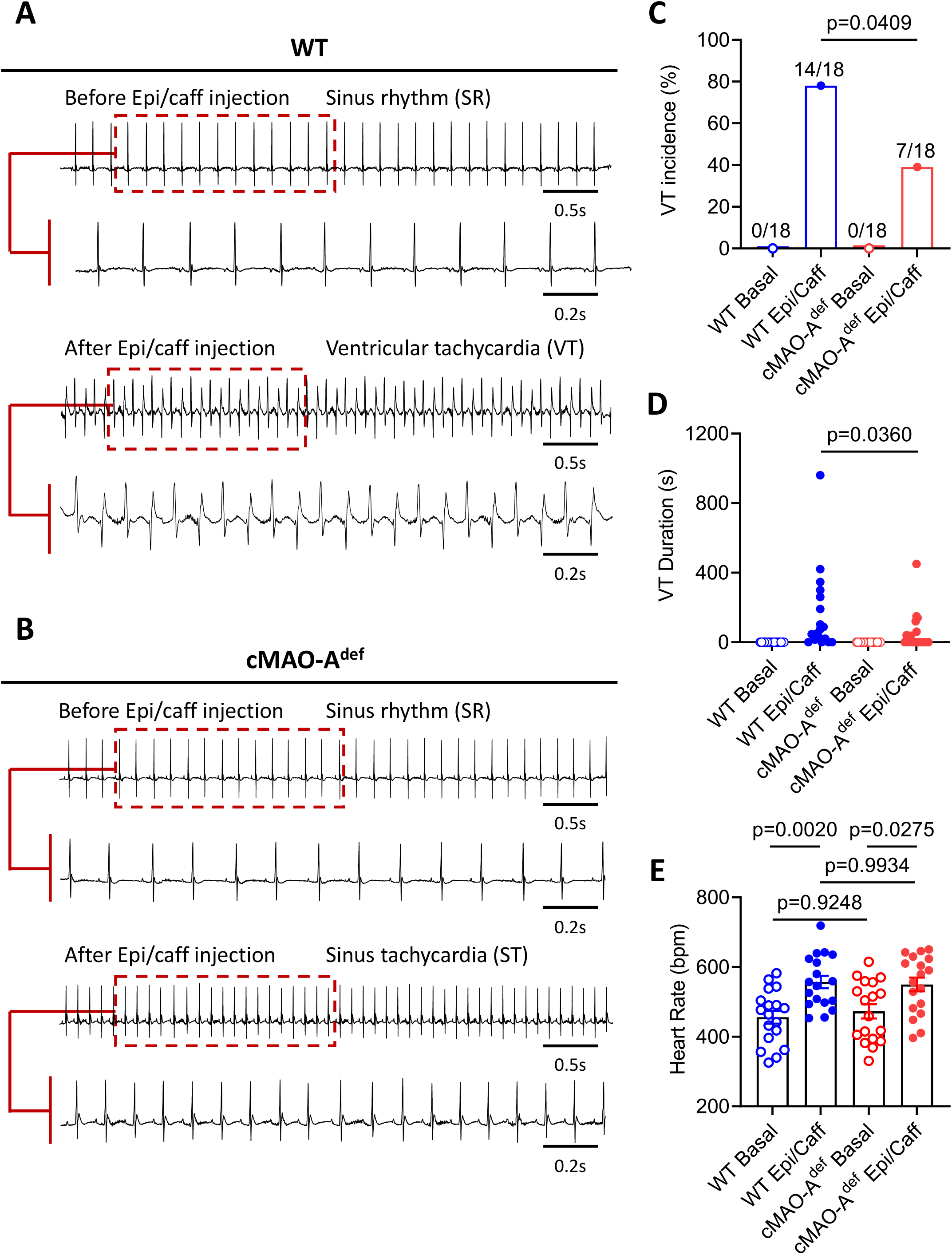
Cardiac MAO-A deficiency blunts susceptibility to catecholamine-induced ventricular tachyarrhythmia (VT) *in vivo*. Representative surface ECG traces obtained from anesthetized WT (A) and cMAO-A^def^ (B) mice before (basal) and after the injection of caffeine (120 mg/kg) and epinephrine (2 mg/kg) to induce VTs. During the 30-min period of ECG recordings following injection, VT incidence (%) (C), VT duration (seconds) (D) and heart rate (bpm) (E) was determined and compared with basal levels of these parameters, in WT or cMAO-A^def^ mice (C-E, N=18 mice per genotype). Data are represented as mean ± SEM in (E). Data are analyzed by Fisher’s exact test in (C), Mann-Whitney test in (D), Paired t-test in (E) for comparison in same genotype; unpaired t-test for comparison between genotypes. P-values of comparisons are shown in the graph.

**Figure 3.**
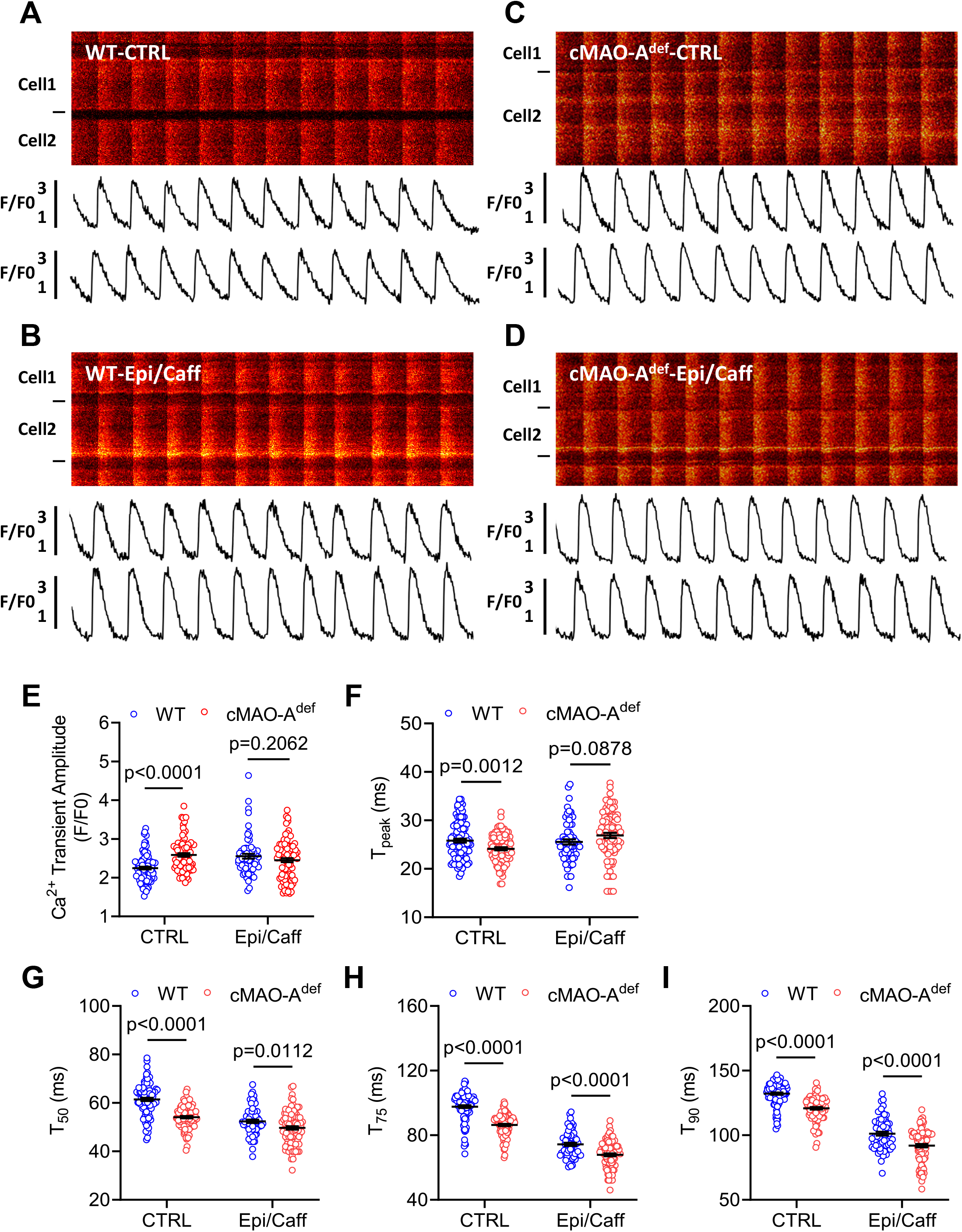
Catecholamine stimulation leads to altered *in situ* Ca^2+^ dynamics in cMAO-A^def^ hearts with 5 Hz pacing. Representative confocal microscopy images of *in situ* Ca^2+^ dynamics driven by 5 Hz electrical stimulation are shown in WT (A, B) and cMAO-A^def^ (C, D) intact hearts at baseline (A, C) and under caffeine (120 μg/ml) and epinephrine (2 μg/ml) perfusion (B, D). The Ca^2+^ transient amplitude (E), time to peak (T_peak_, F), and decay T_50_, (G); T_75_, (H); T_90_, (I) are also shown. (N=67–99 cells from 3-4 hearts/genotype for each group). Cell boundaries were indicated by the black bars on the left. The F/F0 traces depict the average fluorescence signal of the scan area. Data are represented as mean ± SEM. Data are analyzed by unpaired t-test for comparison between genotypes. P-values of comparisons are shown in the graph.

Together, these data demonstrate that cardiac MAO-A inhibition diminishes the susceptibility of the heart to experimental catecholamine stress-induced VT *in vivo*.

### CARDIAC MAO-A INHIBITION ENHANCES DIASTOLIC CA^2+^ HANDLING KINETICS UNDER CATECHOLAMINE STIMULATION IN THE HEART

Imbalanced cellular Ca^2+^ homeostasis underlines arrhythmogenesis^29,30^. Our *in vivo* findings that cardiac MAO-A inhibition suppresses catecholamine induced VT *in vivo* suggests that cardiac MAO-A may affect intracellular Ca^2+^ homeostasis. To this end, we first performed *in situ* Ca^2+^ imaging of ventricular cardiomyocytes from intact mouse hearts attached to an oxygenated Langendorff perfusion system^19^. We examined Ca^2+^ signals initiated by 5 Hz external electric stimulation. At baseline, cardiomyocytes from both WT and cMAO-A^def^ hearts displayed uniform, synchronized Ca^2+^ transients (Figure 3A and 3C). Compared with WT, cMAO-A^def^ cardiomyocytes showed higher Ca^2+^ transient amplitudes (Figure 3E), faster rise times (T_peak_) (Figure 3F) as well as faster Ca^2+^ decay kinetics (Figure 3G-3I), indicating cardiac MAO-A inhibition is associated with accelerated basal Ca^2+^ kinetics.

To understand Ca^2+^ performance in cMAO-A^def^ cardiomyocytes under catecholamine stress, we next perfused 5 Hz stimulated hearts perfused with epinephrine (2 μg/mL) and caffeine (120 μg/mL) (Figure 3B and 3D). Ca^2+^ transient decay kinetics of cMAO-A^def^ cardiomyocytes (mainly reflecting SR Ca^2+^ uptake) were further accelerated beyond those achieved in WT cells (Figure 3G-3I), but without a further increase in Ca^2+^ transient amplitude (Figure 3E). Moreover, sustained catecholamine stimulation under high frequency (50 Hz) stimulation caused aberrant conduction and Ca^2+^ transients in WT hearts (Supplemental Figure 4A). In sharp contrast, cMAO-A^def^ hearts maintained regular electrical patterns and Ca^2+^ dynamics (Supplemental Figure 4B).

**Figure 4.**
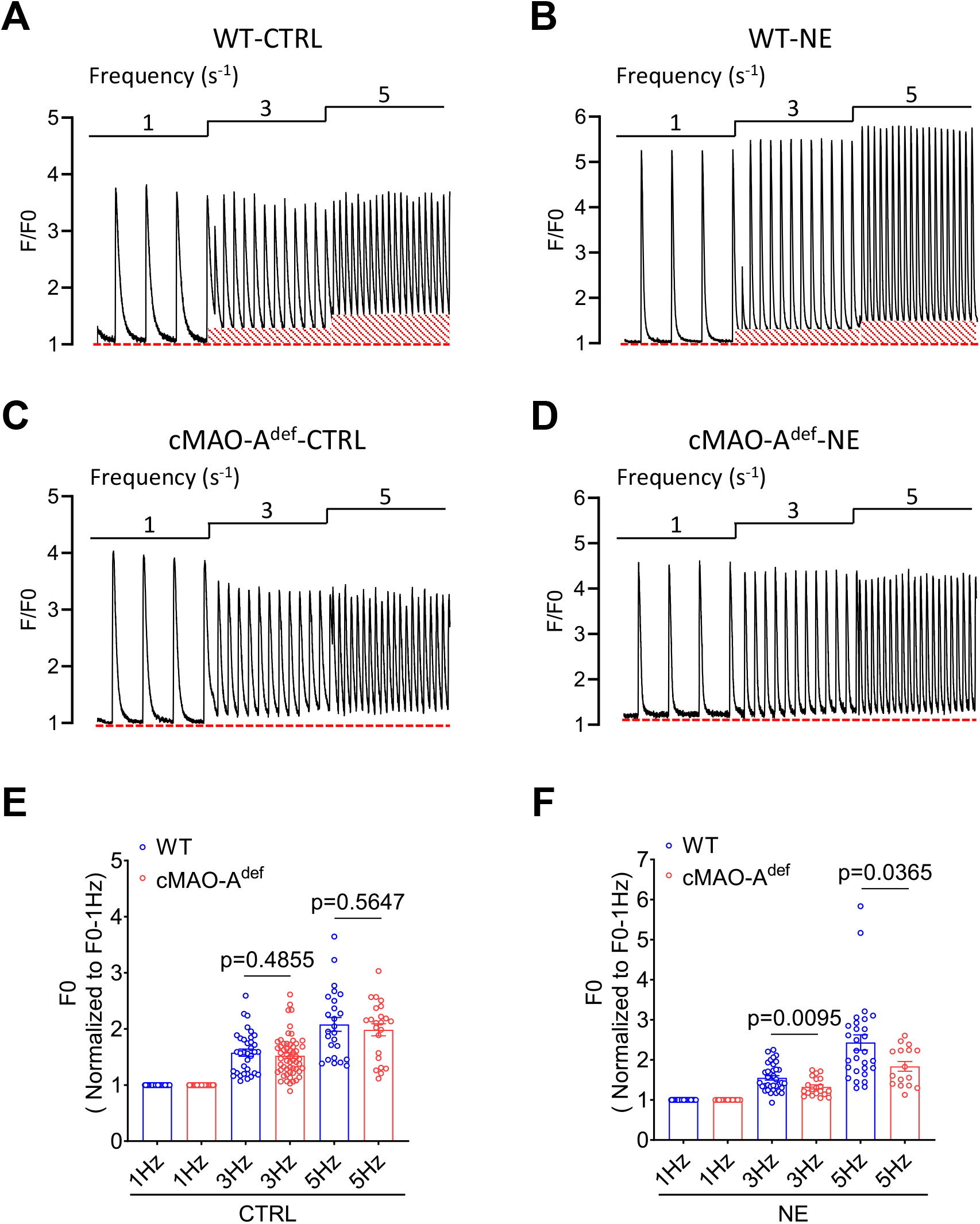
cMAO-A_def_ cardiomyocytes have lower diastolic Ca^2+^ level after catecholamine stimulation. Representative Ca^2+^ transient tracings obtained during 3 different electrical pacing frequencies at basal or under norepinephrine (NE, 1 μM) stimulation in a single cardiomyocyte isolated from WT (A, B) and cMAO-A^def^ (C, D) mouse hearts are shown, highlighting the increased diastolic Ca^2+^ (red box) at higher pacing frequencies in the WT cardiomyocytes. Normalized diastolic Ca^2+^ levels (F0) of WT and cMAO-A^def^ cardiomyocytes at baseline (E) and under NE treatment (F) are also shown. (n=16–57 cells from 3-4 hearts/genotype for each group). Data are shown as mean ± SEM. Data are analyzed by unpaired t-test for comparison between genotypes. P-values of comparisons are shown in the graph.

Taken together, these data suggest that catecholamine stress-induced Ca^2+^ homeostasis and arrhythmias are sensitive to the level of MAO-A activity in the heart.

### CARDIAC MAO-A INHIBITION IMPROVES DIASTOLIC CA^2+^ CONTROL UNDER CATECHOLAMINE STIMULATION IN SINGLE CARDIOMYOCYTES

The decay of the Ca^2+^ transient is largely due to SERCA-mediated reuptake into the SR in rodents^31^. The rate of this decay would be expected to affect end diastolic Ca^2+^ concentration^32^. The faster Ca^2+^ decay kinetics observed in the cardiac MAO-A deficient hearts therefore suggest lower diastolic Ca^2+^ levels. To this end, intracellular Ca^2+^ measurements were performed in isolated cardiomyocytes during field stimulation at 3 different electric frequencies (1, 3, and 5 Hz). Cardiomyocytes from WT hearts showed pacing-induced increase in diastolic Ca^2+^ levels in a frequency dependent manner at baseline (Figure 4A) and under catecholamine stimulation (Figure 4B, norepinephrine, NE). However, the frequency dependent increase in diastolic Ca^2+^ levels was markedly decreased in cMAO-A^def^ cardiomyocytes in response to NE compared to WT cardiomyocytes (Figure 4C-4F).

At the single cardiomyocyte level, studies have shown that higher diastolic Ca^2+^ levels could arise from increased diastolic SR Ca^2+^ leak which in turn activates NCX leading to delayed afterdepolarizations (DADs) and the triggering of VT^33^. Increased sympathetic drive is known to amplify diastolic SR Ca^2+^ leak which can be reflected in increased Ca^2+^ spark activity^34,35^. We therefore measured Ca^2+^ sparks in quiescent cardiomyocytes isolated from WT and cMAO-A^def^ hearts. At baseline, the parameters of Ca^2+^ spark characteristics were comparable between genotypes, including frequency, amplitude, and full duration at half-maximum (FDHM), (Figure 5A-5D). However, in the presence of 1μM norepinephrine (NE), Ca^2+^ spark frequency, amplitude, and FDHM were significantly lower in cMAO-A^def^ cardiomyocytes than WT (Figure 5B-D), suggesting lower Ca^2+^ spark activities in cMAO-A^def^ cardiomyocytes under NE stimulation.

**Figure 5.**
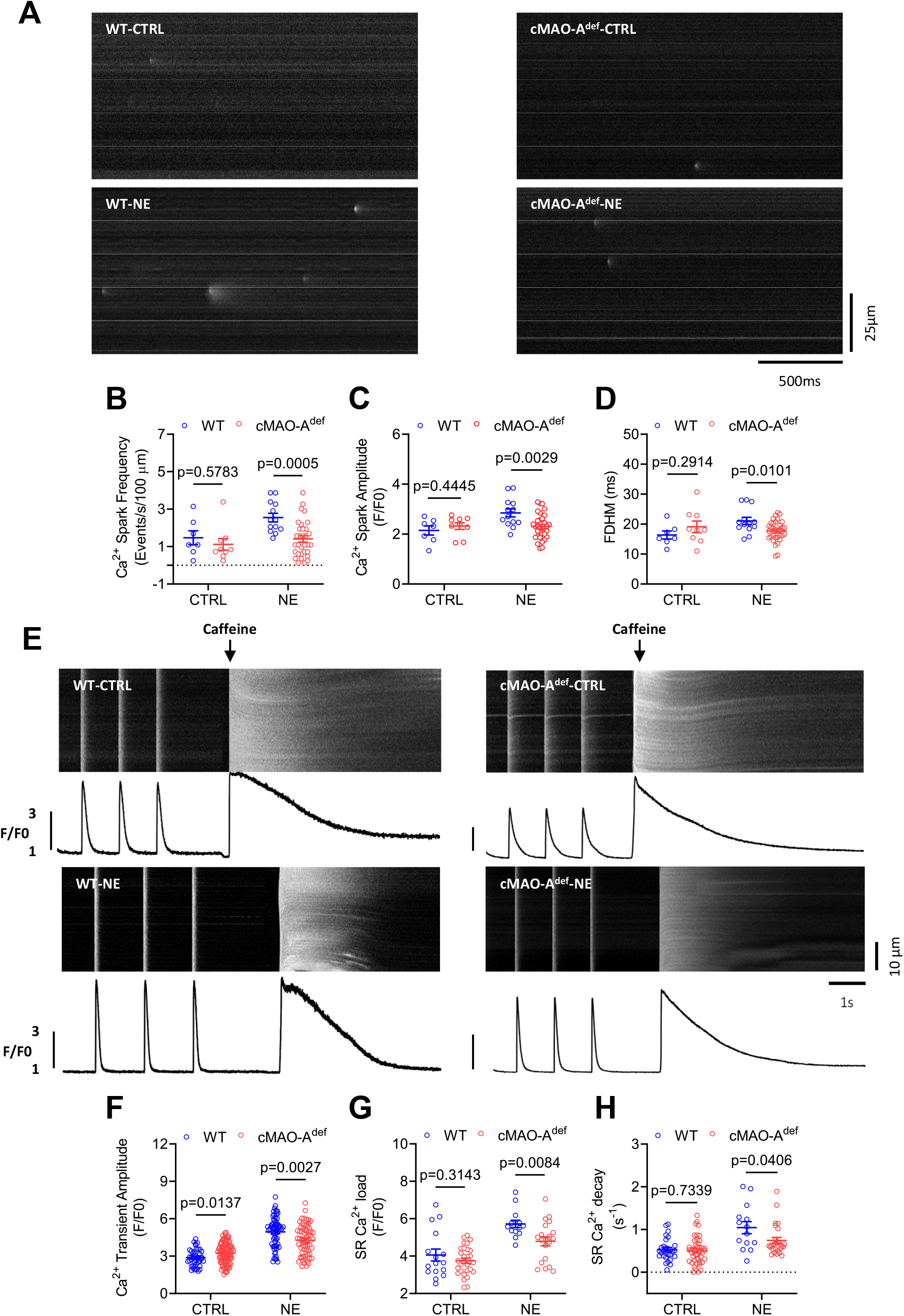
Decreased Ca^2+^ spark activity and reduced sarcoplasmic reticulum (SR) Ca^2+^ content after catecholamine stimulation in cMAO-A^def^ cardiomyocytes. Shown in (A) are representative Ca^2+^ spark images in cardiomyocytes from WT and cMAO-A^def^ mice at baseline or under NE (1 μM) stimulation. The frequency (B), amplitude (C), FDHM (D) of Ca^2+^ sparks are also shown. (n=7-32 cells from 3-4 hearts/genotype per group). SR Ca^2+^ content was measured by rapid caffeine application induced Ca^2+^ release. Shown in (E) are representative traces of 1 Hz field stimulation-triggered Ca^2+^ transients and caffeine-induced Ca^2+^ release (SR Ca^2+^ content) from WT and cMAO-A^def^ cardiomyocytes at basal or under NE stimulation. The amplitude of twitch-induced Ca^2+^ transient at 1Hz electrical stimulation (F), the amplitudes (G) and decay (H) of caffeine-induced SR Ca^2+^ transients are shown (n=16–95 cells from 3-4 hearts/genotype for each group). Data are shown as mean ± SEM. Data are analyzed by unpaired t-test for comparison between genotypes. P-values of comparisons are shown in the graph. FDHM, mean duration at half-peak amplitude; FWHM, full width at half-maximum.

Ca^2+^ sparks are the elementary events of Ca^2+^ release from the SR^36^, and largely affected by the SR Ca^2+^ load^37,38^. We therefore tested whether SR Ca^2+^ content (SR Ca^2+^ load) is altered by measuring the rise in the Ca^2+^ transient induced by rapid application of 20 mmol/L caffeine after a 10 s conditioning electric stimulation at 1 Hz (Figure 5E and 5F). In unchallenged myocytes, SR Ca^2+^ load was comparable between both genotypes. In the presence of 1μM NE, however, SR Ca^2+^ content was significantly lower in cMAO-A^def^ cardiomyocytes compared to those isolated from WT hearts (Figure 5G).

Notably, similar to the *in situ* Ca^2+^ imaging experiments in intact hearts (Figure 3E), twitch-induced Ca^2+^ transient amplitude in isolated cMAO-A^def^ cardiomyocytes was elevated at baseline compared to WT cells (Figure 5F). Under 1μM NE stimulation, however, twitch-induced Ca^2+^ transient amplitude was relatively lower in cMAO-A^def^ cardiomyocytes. The decrease in SR Ca^2+^ content (Figure 5G) and the reduction of twitch-induced Ca^2+^ transient amplitude under NE stimulation (Figure 5F) are consistent and expected because of the known steep relationship that exists between SR Ca^2+^ content and fractional release^39^.

Meanwhile, cMAO-A^def^ cardiomyocytes also showed a slower decay rate of caffeine-induced Ca^2+^ transients under NE stimulation, suggesting Ca^2+^ extrusion via the NCX is slower in cMAO-A^def^ cardiomyocytes (Figure 5H). Since NCX activity has a linear relationship with intracellular Ca^2+^ concentration^40^, slower NCX activity under NE stimulation would be consistent with smaller twitch-induced Ca^2+^ transients observed in cMAO-A^def^ cardiomyocytes (Figure 5F).

Collectively, these data suggest that cardiac MAO-A inhibition is associated with enhanced diastolic Ca^2+^ control in single cardiomyocytes and favors less irregular Ca^2+^ events under catecholamine stress.

### CARDIAC MAO-A INHIBITION INDUCES THE PHOSPHORYLATION OF IMPORTANT CA^2+^ HANDLING PROTEINS IN RESPONSE TO CATECHOLAMINE STIMULATION

We next sought to investigate the molecular determinants of altered Ca^2+^ handling in cardiac MAO-A deficient hearts. Hearts from WT and cMAO-A^def^ mice that underwent baseline recording and catecholamine stress experiments were processed for Western blot studies. First, we inspected expression levels of proteins involving β-adrenergic signaling (a major catecholamine target pathway) and Ca^2+^ handling pathways and observed no significant differences in the protein abundance of β1AR, β2AR, PDE4D, Ca_V_1.2, Na_V_1.5, NCX1, PLM, SERCA2a and CSQ between WT and cMAO-A^def^ hearts at baseline (Supplementary Figure 5).

SR Ca^2+^ reuptake rate is known to be mainly regulated by the SR Ca^2+^ pump SERCA2 and its endogenous inhibitor phospholamban (PLB) during diastole^41,42^ and their regulation could explain the accelerated Ca^2+^ reuptake rate under catecholamine stimulation we observed in cMAO-A^def^ hearts (Figure 3G-3I) Although total SERCA2a and PLB levels are unchanged in WT and cMAO-A^def^ hearts (Supplementary Figure 5 and Figure 6A-6B), the phosphorylation of PLB monomers and pentamers at Serine 16 site was significantly increased after catecholamine stimulation in cMAO-A^def^ hearts (Figure 6C-6D). Serine16 of PLB is phosphorylated by PKA and reflects reduced inhibition of SERCA2 activity^43,44^, which is in line with the faster Ca^2+^ reuptake rates in both *ex vivo* hearts and reduced diastolic Ca^2+^ levels we found in cMAO-A^def^ cardiomyocytes (Figure 3G-3I, Figure 4F). Concurrently, phosphorylation of PLB at threonine 17, a CaMKII or AKT site^45,46^, is unaltered (Figure 6C-6D).

**Figure 6.**
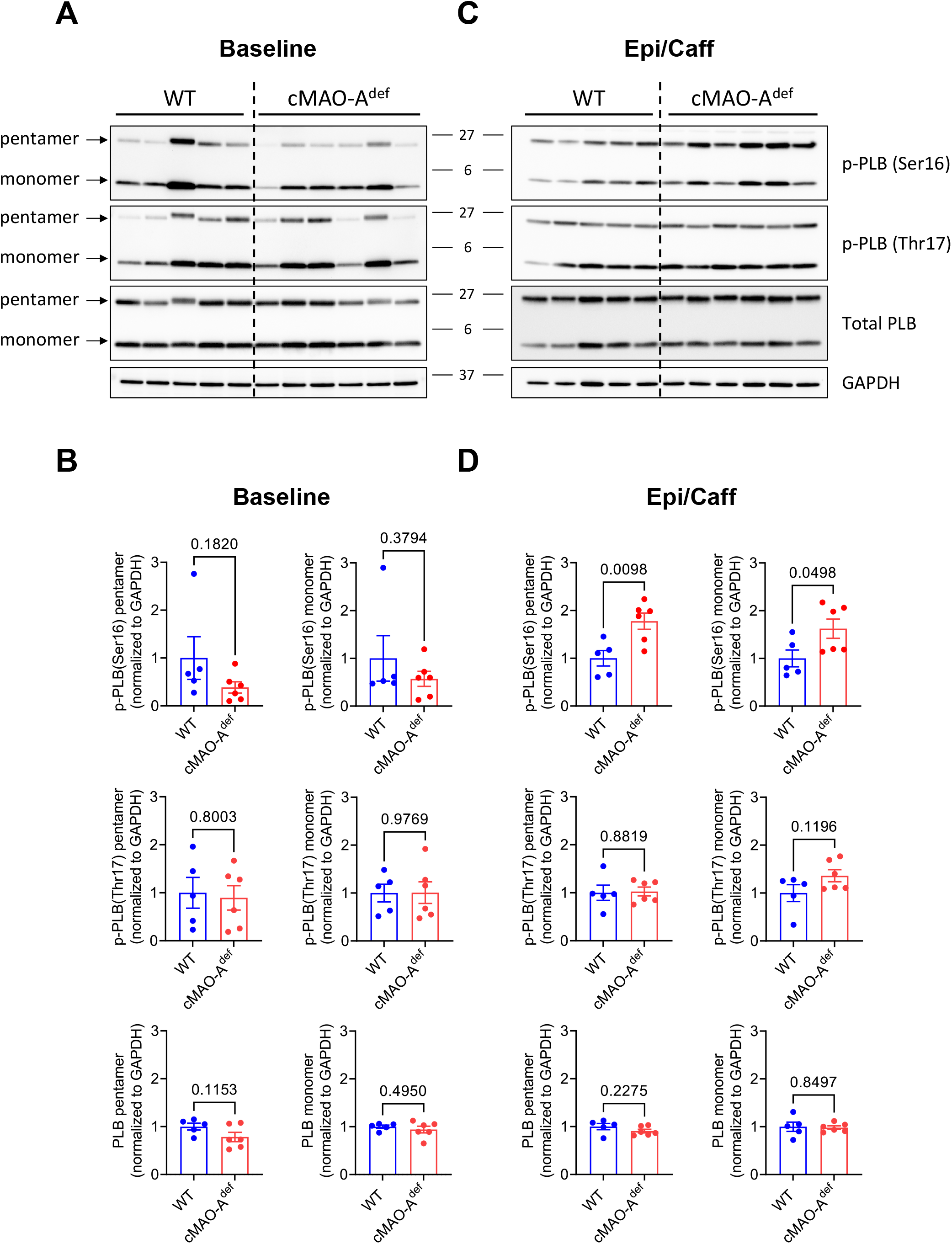
PLB phosphorylation in WT and cMAO-A^def^ myocardium after catecholamine stimulation. Myocardial tissue from WT and cMAO-A^def^ mouse hearts were rapidly collected and frozen in liquid N_2_ at baseline or 30 minutes after injection with caffeine (120 mg/kg) and epinephrine (2 mg/kg). Representative immunoblots of p-PLB (Ser-16), p-PLB (Thr-17) and total PLB at baseline (**A**) and 30 minutes after epinephrine/caffeine challenge (**C**) are shown (N=5-6 mice for each genotype). Densitometric quantification of total and phosphorylated levels of the PLB pentamer and monomer at baseline (**B**) and after epinephrine/caffeine (**D**). Data are normalized to GAPDH and shown as mean ± SEM. Data are analyzed by unpaired t-test. P-values of comparisons are shown in the graph. PLB, phospholamban.

Lower diastolic Ca^2+^ levels could be both a cause and an effect of reduced diastolic SR Ca^2+^ leak and is supported by our observations that cMAO-A^def^ cardiomyocytes have reduced Ca^2+^ spark amplitude and frequency under catecholamine stimulation (Figure 5A-5D). Ca^2+^ release from the SR is mediated through RyR2 channels and increased through post translational modification^33,47,48^. In support of this hypothesis, we found a significant increase in RyR2 phosphorylation at serine 2814 in cMAO-A^def^ hearts at baseline, but not serine 2808 and serine 2030 (Figure 7A-7B). Serine 2814 phosphorylation is likely to have contributed to the larger Ca^2+^ transients observed in cMAO-A^def^ cardiomyocytes at baseline (Figure 3E and Figure 5F). In sharp contrast, serine 2814 phosphorylation is significantly reduced after catecholamine stress (Figure 7C-7D). Serine 2814 of RyR2 is a well-known target of CaMKII, which has a prominent role in the pathophysiology of both heart failure and arrhythmias ^33,47,48^. Consistently, CaMKII autophosphorylation at threonine 286 site, a marker of CaMKII activation, was significantly suppressed in cMAO-A^def^ hearts after catecholamine stress relative to WT mice (Figure 7C-7D).

**Figure 7.**
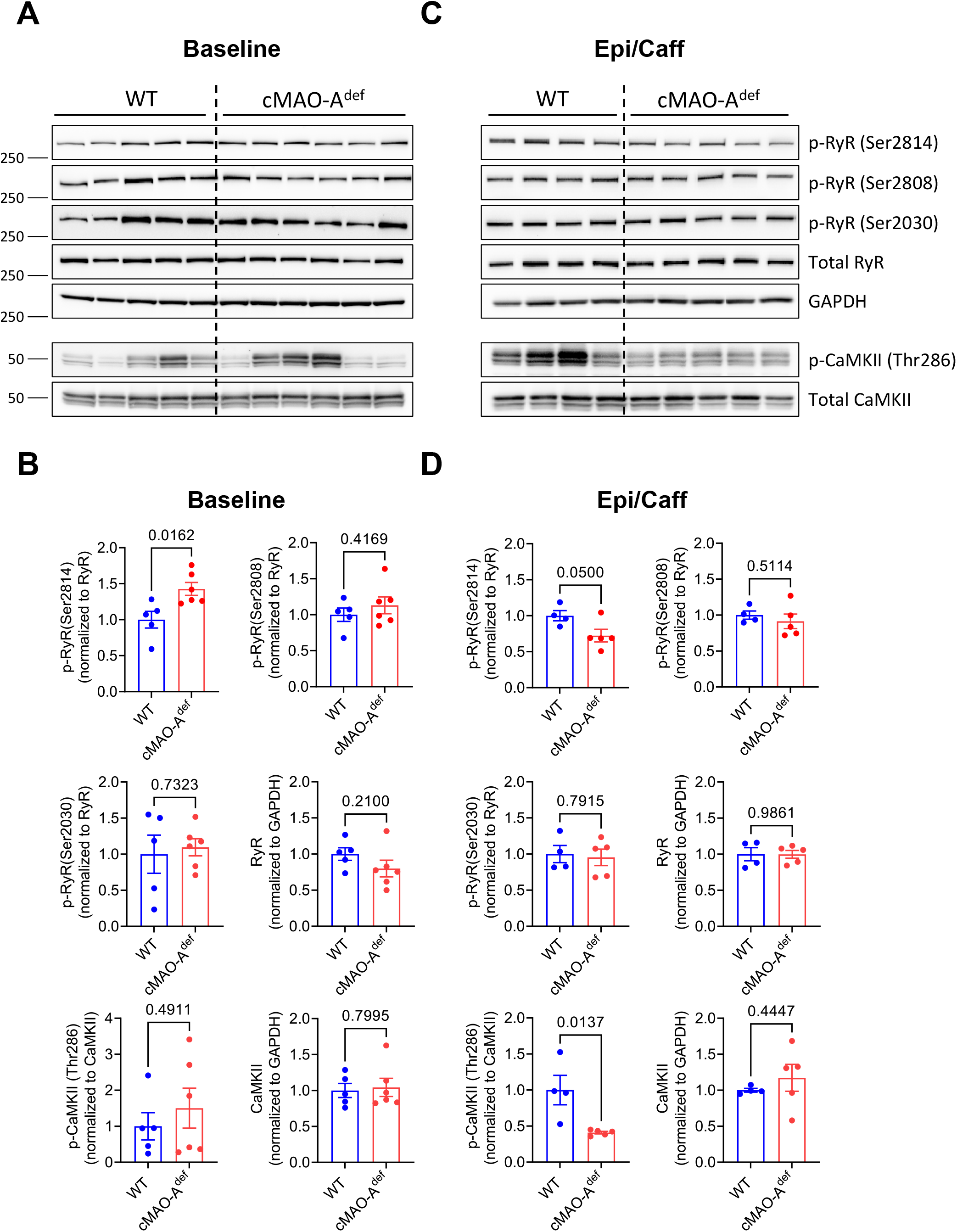
CaMKII and RyR phosphorylation in WT and cMAO-A^def^ myocardium after catecholamine stimulation. Representative immunoblots of p-RyR2 (Ser-2814), p-RyR2 (Ser-2808), p-RyR2 (Ser-2030), p-CaMKII (Thr-286), total RyR2 and CaMKII at baseline (**A**) and 30 minutes after epinephrine/caffeine challenge (**C**) are shown (N=5-6 mice for each genotype). Densitometric quantification of total and phosphorylated RyR2 and CaMKII at baseline (**B**) and 30 minutes after epinephrine/caffeine challenge (**D**). Total RyR2 or CaMKII protein is normalized to GAPDH. Phosphorylated RyR2 and CaMKII are normalized to corresponding total protein. Data are represented as mean ± SEM. Data are analyzed by unpaired t-test. P-values of comparisons are shown in the graph.

Together, these data suggest that cardiac MAO-A inhibition alters the phosphorylation status of Ca^2+^ handling proteins leading to accelerated SR Ca^2+^ uptake but limiting spontaneous SR Ca^2+^ release upon stimulation that involves the suppression of CaMKII activation.

## DISCUSSION

Arrhythmogenesis is a complex and multifactorial cardiac disorder involving electrical and metabolic imbalances which serve as the substrate for an initiating trigger. Catecholaminergic stress is a well-known arrhythmogenic trigger. The present study provides clinical and experimental evidence that MAO-mediated catecholamine metabolism can contribute to arrhythmia induction. We found that cardiac MAO-A inhibition is significantly associated with reduced arrhythmic events in humans and diminishes catecholamine triggered arrhythmic events in mice. At the cellular level, our findings clearly show that in response to catecholamine stimulation, cMAO-A^def^ hearts have faster rates of Ca^2+^ reuptake that correlates with increased PLB phosphorylation at its PKA site (serine 16), resulting in lower diastolic Ca^2+^ levels. Concurrently, we observed decreased CaMKII activity in the cMAO-A^def^ hearts reflected by significantly reduced CaMKII autophosphorylation at its threonine 286 site. One important downstream target of CaMKII is RyR2. CaMKII dependent phosphorylation of RyR2 promotes diastolic Ca^2+^ leak in the form of Ca^2+^ sparks that ultimately leads to arrhythmic Ca^2+^ waves^34^. Consistent with lower CaMKII phosphorylation, RyR2 phosphorylation at its CaMKII site (serine 2814) is significantly reduced in cMAO-A^def^ hearts after catecholamine stimulation. Accordingly, diastolic Ca^2+^ spark activity is also dramatically reduced. Together, these findings suggest that MAO-A inhibition in the heart results in improved diastolic Ca^2+^ homeostasis that protects against catecholamine related arrhythmic triggers. Our results therefore implicate cardiac MAO-A as a targetable candidate for preventing arrhythmogenesis.

Depression is often associated with aberrant sympathoadrenal activity ^49–53^, representing a clear pathophysiologic pathway through which these disorders increase cardiovascular disease risk^49,50^. Of note, depression is associated with a substantially increased risk of atrial fibrillation and ventricular arrhythmia^2–4^. Even though MAO inhibitors have been prescribed for decades for the treatment of depression^5^, detailed mechanistic analysis of the effects of antidepressant therapy on adverse arrhythmic outcomes in depressed patients has not been well documented. Our findings, based on a large and well-balanced clinical cohort suggest that patients with a diagnosis of depression and treated with MAOI had lower risk and encountered less incidents of adverse arrhythmic outcomes compared to those treated with SSRI. Although the effect size is small, likely due to numerous cardiometabolic factors which cannot be accounted for, the difference is significant and could be of interest to cardiologists (Figure 1). MAO inhibitors are currently no longer the first line choice of antidepressants, mainly because MAO isoforms are expressed throughout the body and their inhibition produces undesirable side effects^54^. Our analysis of clinic databases, however, reveals a potential anti-arrhythmic effect of MAO inhibitors that warrant the generation of cardiac specific MAO inhibitors for clinical use in patients with cardiac arrhythmias.

Aberrant Ca^2+-^ handling in cardiomyocytes caused by excessive β-adrenergic activation are a leading mechanism linking catecholaminergic overload to arrhythmogenesis^34^. At the level of the cardiomyocyte, spontaneous Ca^2+^ release from hyperactive RyR2 channels during diastole coalesce into a propagated Ca^2+^ wave, which in turn activates the electrogenic NCX, resulting in cell membrane depolarization. Membrane depolarizations of sufficient amplitude will activate voltage-gated Na^+^ channels, which in turn will trigger a full cardiac action potential^55^. The anti-arrhythmic effect in cMAO-A^def^ mice therefore may be mediated by altered β-adrenergic responses and/or Ca^2+^ handling processes under catecholamine stress. Our examination of the β-adrenergic signaling pathway including β1AR, β2AR and PDE4D revealed that no expression changes were observed between WT and cMAO-A^def^ hearts, suggesting no adaptation occurred on the protein expression level of this pathway. However, in line with the recent report that an intracellular pool of β1AR is located at the SR and contributes to the inotropic effect of catecholamines via enhanced cAMP/PKA pathway activity, we find phosphorylation of PLB at serine16 (a PKA target site) was substantially elevated after catecholamine stimulation in cMAO-A^def^ hearts compared to WT hearts. As PLB is the major inhibitory regulator of SERCA2, phosphorylation of PLB during β-adrenergic stimulation relieves its inhibitory effects and promotes Ca^2+^ reuptake into the SR during diastole^42^. Consistent with the higher PLB phosphorylation, faster Ca^2+^ uptake rates and reduced diastolic Ca^2+^ level was observed in cMAO-A^def^ cardiomyocytes (Figure 3G-3I, Figure 4A-4F).

Sustained activation of CaMKII is also known to underlie arrhythmogenesis^56^. CaMKII activation is driven by Ca^2+^ bound calmodulin in response to an increase in cytosolic Ca^2+^ within the cardiomyocyte^57^. Once activated, CaMKII then phosphorylates itself (at threonine 286), thereby prolonging and potentiating its activated state^58^. Consistent with faster Ca^2+^ reuptake rate and lower diastolic Ca^2+^ levels (Figure 3G-3I, Figure 4A-4F), CaMKII auto-phosphorylation was significantly decreased in response to catecholamine stress in cMAO-A^def^ (Figure 7C and 7D), indicating reduced activity. CaMKII targets several proteins involved in the regulation of cardiomyocyte electrophysiology and Ca^2+^ fluxes including RyR2. The significantly reduced CaMKII-dependent phosphorylation of RyR2 (serine 2814) (Figure 7C and 7D) likely contributed to the reduction in amplitude and frequency of spontaneous Ca^2+^ releases during diastole (Figure 5A-5D) in cMAO-A^def^ cardiomyocytes. This observation is consistent with the previous report of transgenic mice with RyR-S2814 (S2814A) ablation in which spontaneous Ca^2+^ waves were reduced, rendering them protected from catecholaminergic-induced arrhythmias^59^.

In addition, CaMKII can also be activated by oxidative stress and contribute to arrhythmias^60^. Previous work from our group and others have shown that cardiac MAO-A are robust generators of H_2_O_2_ and reactive catecholaldehydes^12,61^, the latter of which causes disruptions in mitochondrial oxidative phosphorylation (OxPHOS)^62^, and stimulates pro-inflammatory/pro-fibrotic signaling in the myocardium^63^. A decrease in oxidative stress in cMAO-A^def^ hearts following catecholaminergic overload would thus predictably avoid the overaction of CaMKII and other adverse effects, all of which could contribute to the reduced arrhythmogenesis found in the cMAO-A^def^ mice. This anti-arrhythmic effect in the cMAO-A^def^ mice would be potentially beneficial in the context of aging, diabetes and heart failure, conditions which are associated with increased cardiomyocyte MAO expression in parallel with risk of arrhythmias ^12,13,62,64,65^. Other targets of CaMKII in cardiomyocytes such as Na^+^ channels (Na_V_1.5) and L-Type Ca^2+^ channels (LTCC), which are also known to be involved in arrhythmogenesis (reviewed in ^66,67^), could also be potentially involved.

## CONCLUSIONS

Taken together, our data provide evidence for the anti-arrhythmic effect of cardiac MAO-A inhibition, which mechanistically is linked with enhanced Ca^2+^ reuptake to lower diastolic Ca^2+^ level in cardiomyocytes. Our findings further implicate the therapeutic potential of targeting cardiac MAO-A for prevention and treatment of cardiac arrhythmias.

---

**Supplementary Figure 1.**
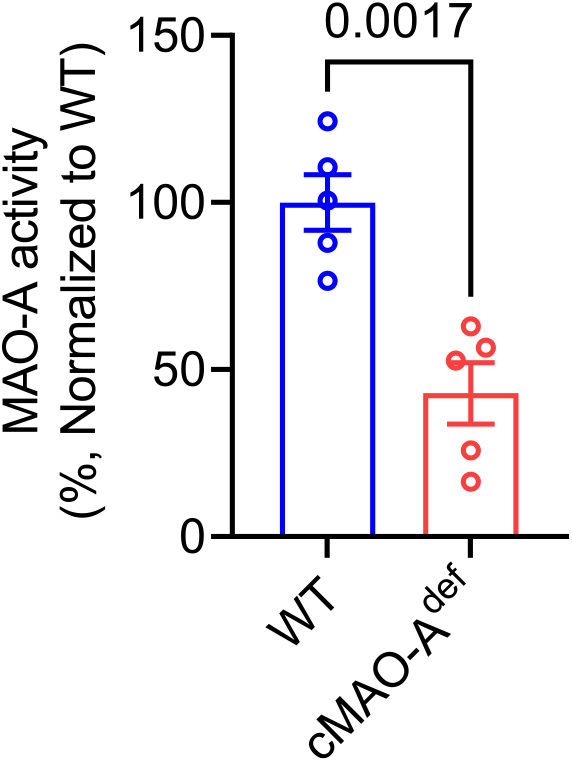
MAO-A activity in isolated cardiomyocytes prepared from WT and cMAO-A^def^ heart. MAO-A activity is decreased in isolated cardiomyocytes from cMAO-A^def^ mouse heart (N=5 mice per genotype, aged 2–3-month-old). Data are shown as mean ± SEM. Data are analyzed by unpaired t-test. P-values are shown in the graph.

**Supplementary Figure 2.**
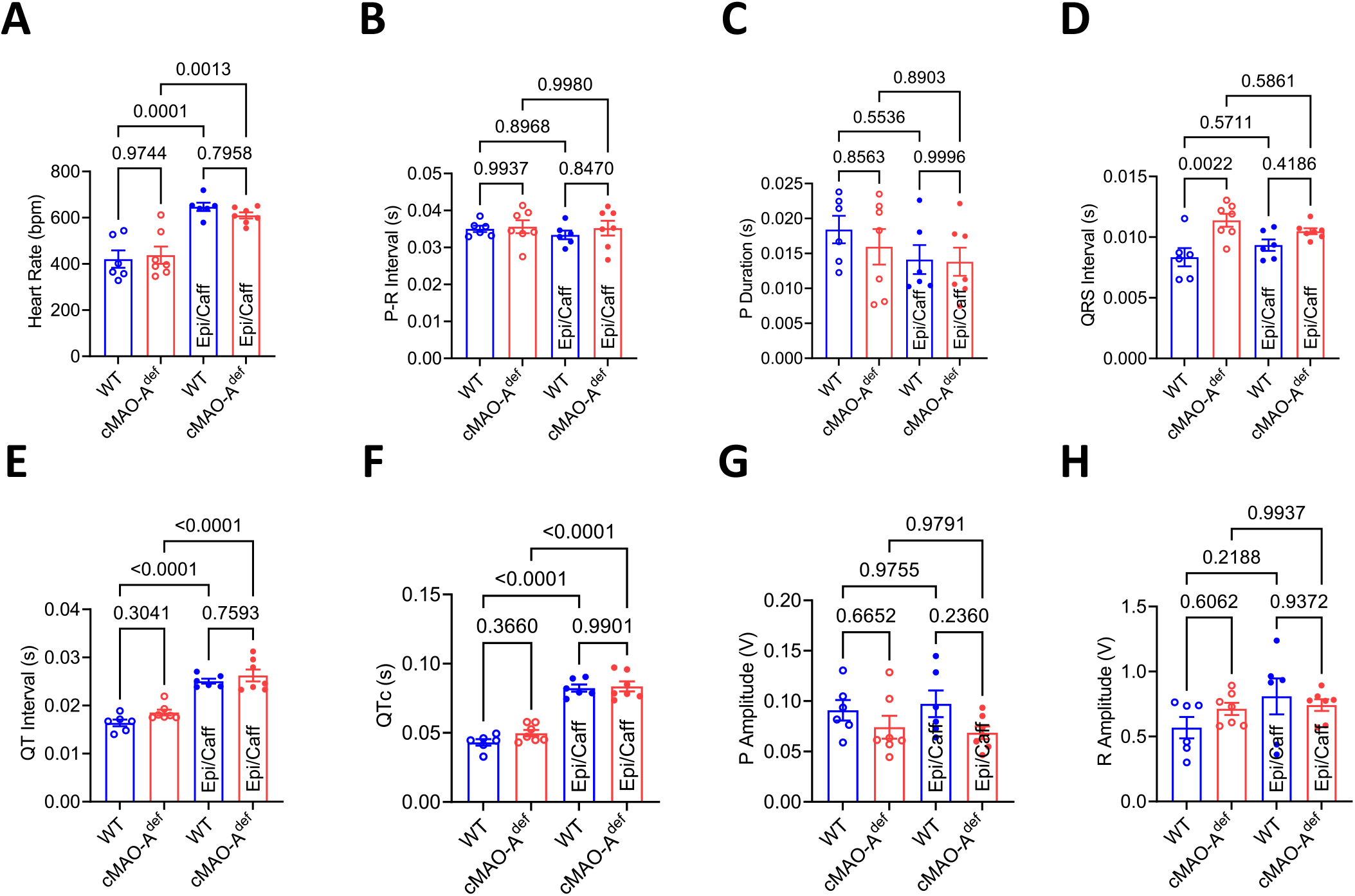
Surface ECG parameters of WT and cMAO-A^def^ mice. Quantification of the indicated ECG parameters from WT and cMAO-A^def^ mice before and at the end of 30 min injection of caffeine (120 mg/kg) and epinephrine (2 mg/kg) challenge (N=6-7 mice per genotype). Data are represented as mean ± SEM. Data are analyzed by one-way ANOVA followed by Tukey’s multiple comparison test. P-values of comparisons are shown in the graph.

**Supplementary Figure 3.**
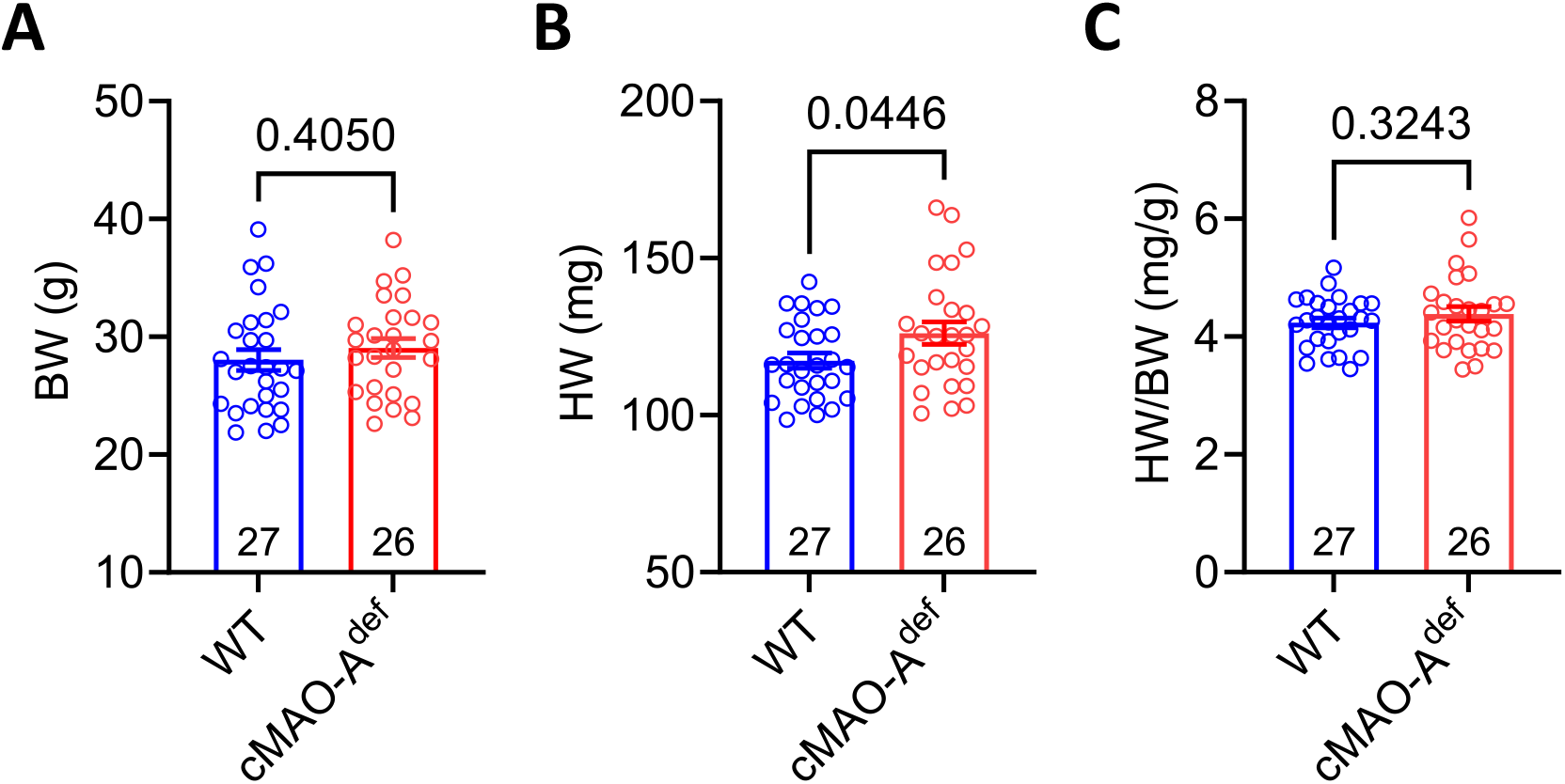
Body weight (BW) and heart weight (HW) of WT and cMAO-A^def^ mice. BW (A), HW (B) and HW/BW (C) of WT and cMAO-A^def^ male mice are shown at 2-3 months of age. Numbers of mice for each genotype used for this analysis are indicated inside the bars of the graph. Data are represented as mean ± SEM. Data are analyzed by unpaired t-test. P-values of comparisons are shown in the graph.

**Supplementary Figure 4.**
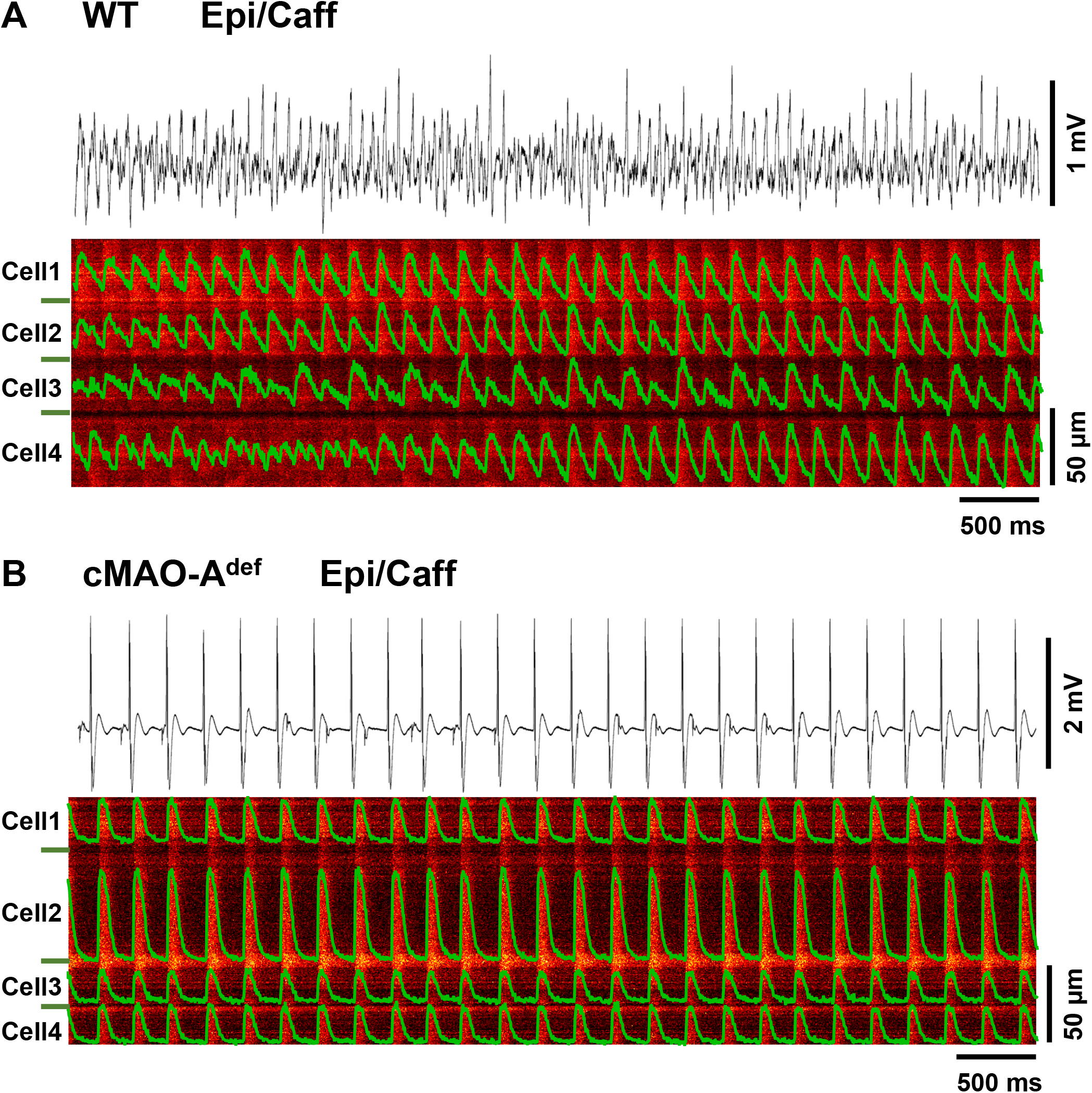
*In situ* confocal Ca^2+^ imaging and ECGs of intact WT and cMAO-A^def^ hearts after 50 Hz pacing under catecholamine stimulation. Representative Ca^2+^ transients after 50 Hz electrical stimulation in WT (A) and cMAO-A^def^ (B) intact hearts under caffeine (10 mg/ml) and epinephrine (1.6 mg/ml) perfusion. Cell boundaries are indicated by the green bars on the left. The green F/F0 traces depict the average fluorescence signal of the scan area. The corresponding ECG recordings are shown above the Ca^2+^ transients traces.

**Supplementary Figure 5.**
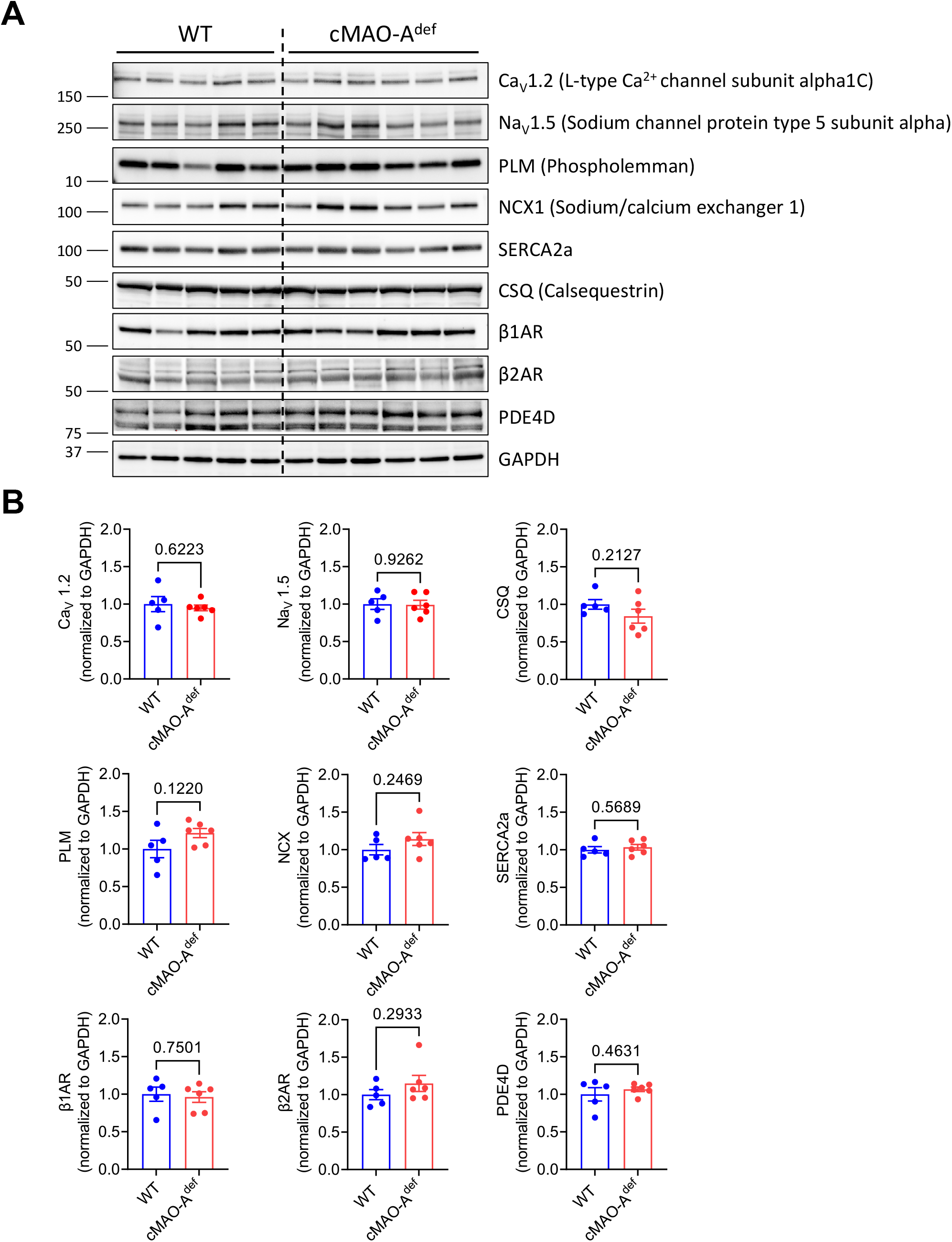
β-adrenergic pathway and Ca^2+^ handling protein expression in unstimulated WT and cMAO-A^def^ mouse hearts. Shown in (A) are representative immunoblots of proteins involved in β-adrenergic signaling pathway and Ca^2+^ handling detected by corresponding antibodies as indicated (N=5-6 mice for each genotype), along with the corresponding densitometry analysis (B). Relative expression of proteins of interest are normalized to GAPDH and shown as mean ± SEM. Data are analyzed by unpaired t-test. P-value of comparisons are shown in the graph.

